# Designing Bt constructs for Brassicas, with minimal IP issues – A case study

**DOI:** 10.1101/2024.01.09.574921

**Authors:** Md Mahmudul Hassan, Francis Tenazas, Adam Williams, Jing-wen Chiu, Charles Robin, Derek A. Russell, John F. Golz

**Affiliations:** School of Biosciences, University of Melbourne, Parkville, VIC 3010, Australia; Department of Genetics and Plant Breeding, Patuakhali Science and Technology University, Dumki, Patuakhali-8602, Bangladesh; School of Agriculture, Food and Ecosystem Sciences, University of Melbourne, Parkville, VIC 3010, Australia; Veterinary School, University of Melbourne, Parkville, VIC 3010, Australia

**Keywords:** *Bacillus thuringiensis*, insecticidal gene, Bt toxin, gene stacking, Diamondback moth, *Cry1B*, *Cry1C*, *Cry1*^*M*^

## Abstract

As part of a publicly funded initiative to develop genetically engineered Brassicas (cabbage, cauliflower, and canola) expressing *Bacillus thuringiensis Cry*-encoded insecticidal (Bt) toxin for Indian and Australian farmers, we designed several constructs that drive high-level expression of modified *Cry1B* and *Cry1C* genes (referred to as *Cry1B*^*M*^ and *Cry1C*^*M*^). The two main motivations for modifying the DNA sequences of these genes were to minimise any licencing cost associated with the commercial cultivation of transgenic crop plants expressing *Cry*^*M*^ genes, and to remove or alter sequences that might affect gene activity in plants. To assess the insecticidal efficacy of the *Cry1B*^*M*^*/Cry1C*^*M*^ genes, constructs were introduced into the model Brassica *Arabidopsis thaliana* in which *Cry1B*^*M*^*/Cry1C*^*M*^ expression was directed from either single (*S4*/*S7*) or double (*S4S4*/*S7S7*) Subterranean Clover Stunt Virus promoters. The resulting transgenic plants displayed a high-level of *Cry1B*^*M*^/*Cry1C*^*M*^ expression. Protein accumulation for *Cry1C*^*M*^ ranged from 0.81 to 17.69 μg Cry1C^M^/g fresh weight of leaves. Contrary to previous work on stunt promoters, we found no correlation between the use of either single or double stunt promoters and the expression levels of *Cry1B*^*M*^*/Cry1C*^*M*^ genes, with a similar range of *Cry1C*^*M*^ transcript abundance and protein content observed from both constructs. First instar Diamondback moth (*Plutella xylostella*) larvae fed on transgenic Arabidopsis leaves expressing the *Cry1B*^*M*^*/Cry1C*^*M*^ genes showed 100% mortality, with a mean leaf damage score on a scale of zero to five of 0.125 for transgenic leaves and 4.2 for wild-type leaves. Under laboratory conditions, even low-level expression of *Cry1B*^*M*^ and *Cry1C*^*M*^ was sufficient to cause insect mortality, suggesting that these modified *Cry*^*M*^ genes are suitable for the development of insect resistant GM crops. Except for the *Cry1B/Cry1C* genes themselves, which remain under patent until 2027 and the *PAT* gene in the USA, our assessment of the intellectual property landscape of the constructs described here suggest that they can be used without the need for further licencing. This has the capacity to significantly reduce the cost of developing and using these *Cry1*^*M*^ genes in GM crop plants in the future.

## 1.0 Introduction

Introducing *Crystal* (*Cry*) genes from the soil bacteria *Bacillus thuringiensis* into commercially grown crop plants is a highly effective strategy to control insect pests, as insects across broad taxonomic groupings are susceptible to the encoded Bt toxins [1]. However, a common problem associated with this control strategy is the development of insect resistance to the Bt toxin present in the transgenic plants [2, 3]. Several approaches have been used to reduce or prevent the development of insect resistance including the use of refuge crops (providing a sufficiently high populations of susceptible insects to prevent resistance genes from becoming homozygous), high expression of *Cry* genes in plants, deploying different *Cry* genes in individual plants in a crop (seed mixtures), and combining multiple *Cry* genes (i.e., stacking) in the same plant [4–8]. Of these, high expression and stacking of *Cry* genes in the same plant are considered the most effective strategies [1, 5, 9, 10]. For example, plants expressing both *Cry1Ac* and *Cry1C* genes greatly delayed the emergence of resistance to the encoded toxins by Diamondback moth (DBM) (*Plutella xylostella*) [11]. Plants with stacked *Cry* genes are also protected from insects that are less susceptible to Bt toxins such as *Helicoverpa armigera* [12]. For this reason, plants harbouring stacked *Cry* genes are favoured by companies developing Bt crops as exemplified by the replacement of GM cotton containing a single *Cry* (*Cry1Ac*) gene with a gene stack (*Cry1Ac*/*Cry2Ab*) [7]. Although plants with stacked *Cry* genes have been successful in controlling insect pests in the field, there is still the potential for resistance to develop The most common form of insect resistance to a Bt toxin is associated with a mutation in the receptor that binds to the toxin in the insect mid-gut [13–15]. Therefore, selection of *Cry* genes used for stacking is an important factor determining durability of the Bt toxin in the field, as different Bt toxins may bind to different receptors with different strengths. As these binding patterns are becoming increasingly well understood, it is now possible to optimize stacking by selecting Bt genes that are not susceptible to known resistance mechanisms in particular insect targets.

High-level accumulation of Bt toxin within plant tissues is generally lethal to insects that are either fully susceptible or have a single copy of a recessive gene for resistance [2, 9, 16, 17]. *Cry* gene expression in plants depends on many factors including their nucleotide structure, the promoter used to drive their expression, and the location and copy number of the *Cry* gene within a plant genome [18]. A suboptimal nucleotide structure is among the main factors contributing to low *Cry* gene expression in plants as, due to their bacterial origin, *Cry* genes contain many sequences that negatively impact on protein production in eukaryotic cells. The presence of signal sequences required for polyadenylation, mRNA decay and splicing, also affects mRNA structure and accumulation in plants [19–21]. For example, the presence of three AATAAA repeats within the coding region of *CryIIIA* is associated with premature polyadenylation of the gene when expressed in plants, as these sequences match the polyadenylation signal usually found in the 3’-untranslated region of many eukaryotic genes [22–24]. In addition, *Cry* gene expression in plants is impacted by differences in nucleotide content between bacterial and eukaryotic genomes. For instance, *Cry* genes have a higher AT content (65%) compared to typical dicot (55%) or monocot (45%) plant genes [25]. These differences mean that the *Cry* genes utilize codons that are less common in plants, which reduces the rate of protein production due to the limited size of tRNA pools for these codons [26]. Furthermore, if a ribosome fails to incorporate the corresponding tRNA for a rare codon, translation may be aborted, resulting in the ribosome becoming disassociated from the mRNA. Poorly translated mRNAs are prone to degradation in the host cell by nonsense mediated RNA decay [20]. Rectifying these issues, together with the removal of spurious polyadenylation signal sequences and sequences that might be responsible for mRNA instability, such as the ATTTA motif, from plant-expressed *Cry* genes can significantly improve production of the encoded Bt toxin in plants [21, 27–30]. For instance, by changing the composition of codons so that they better reflect the distribution of those seen in typical plant genes, significant increases in Bt protein have been observed in transgenic tobacco, tomato and potato plants [21].

Commercialization of GM crops requires the developer to manage several legal hurdles, due to the complex patent landscape associated with the technologies used in the creation of GM crops. Almost all the significant components of the constructs used in plant transformation are patented. These include the ‘effect gene’ and associated regulatory sequences, as well as the selectable marker [31]. For example, use of an antibiotic resistance gene as selectable marker in plant transformation is restricted by a patent owned by Monsanto, however, this IP right only applies in the US. Another example of a patent that has a considerable impact on construct design is the use of the cauliflower mosaic virus (CaMV) *35S* promoter to drive selectable marker gene activity in plants [31]. Patent holders frequently do not allow access to a patented technology if they are themselves using it commercially or have sole licensing agreements with the other entities. Therefore, at early stages of GM crop development, Freedom to Operate (FTO) needs to be established for technologies used in introducing new traits to crops of interest. Without securing all the necessary legal rights, GM crop developers may be exposed to legal liabilities, which ultimately prevent the use of the developed crop. A notable example of the complexity associated with IP issue is the development of golden rice, a transgenic rice line rich in ß-carotene (a precursor of vitamin A). Delivery of the golden rice for public use was delayed due to extensive patenting issues, associated with 72 patents owned by 40 organization [31–33].

As a part of Australian-Indian government strategic initiative, our aim was to develop Bt-expressing Brassica crop for commercial use in both countries where the licencing costs associated with the use of this technology was minimized. Here we describe the generation of a *Cry1B*^*M*^/*Cry1C*^*M*^ gene stack that may be used as an effective insecticide when introduced into plants. Nucleotide modification of the *Cry1B*/*Cry1C* genes, together with careful selection of components used in the design of the constructs, ensured both high-level expression in plants and minimal licensing costs associated with the use of these constructs. We demonstrate under laboratory conditions that Arabidopsis plants expressing the modified *Cry* genes display high-level resistance to diamondback moth (DBM) larvae, consistent with our modifications not adversely affecting the lethality of the *Cry* genes. The results of this study provide an example of how new Bt-expressing constructs that are relatively free of third-party IP may be generated, particularly for deployment in developing nations where farmers may have limited capacity to pay the additional costs associated with Bt crops developed by multinational seed companies.

## 2.0 Materials and Methods

### Plant materials and growth conditions

The Columbia-0 ecotype of *Arabidopsis thaliana* was used in this study. Seeds were grown on a soil/perlite mix or half-strength Murashige and Skoog (½MS) media containing Phytagel. Seeds were stratified at 4°C for 2-3 days prior to placement in a growth room under continuous light at 18-20°C or a growth cabinet under continuous light at 20-22 °C.

### Selection of components for *Cry1B*^*M*^/*Cry1C*^*M*^ constructs

The *Cry1B*^*M*^ construct was designed to have either one or two *S4* subterranean clover stunt virus (SCSV) promoters [34] upstream and the pea *RUBISCO E9* terminator [35] downstream of the *Cry* coding sequence. In contrast, either one or two SCSV *S7* promoters [34] were placed upstream and the *Flaveria bidentis MALIC ENZYME* (*ME*) terminator [34] downstream of the *Cry1C*^*M*^ gene (Figure 1). Our previous work with *Cry1B/Cry1C* genes identified leaky expression of *Cry1C* in *E. coli*. To prevent this, an intron from potato *ST-LS1* gene [36] was placed within the *Cry1C*^*M*^ codign sequence. A DNA fragment containing these elements (*ME*_*ter*_*:Cry1C*^*M*^*-intron::S7S7-S4S4::Cry1B*^*M*^*::E9*_*ter*_) was then synthesized to our specifications by Biomatik (www.biomatik.com) and cloned in the *Eco*RI/*Hin*dIII sites of the pUC19 vector. This vector was subsequently digested with *Bgl*II enzyme to remove one copy of the *S4* and *S7* promoters resulting in single stunt promoter constructs (*ME*_*ter*_ *Cry1C*^*M*^*-intron::S7-S4::Cry1B*^*M*^*::E9*_*ter*_*). Cry1*^*M*^ genes under single or double stunt promoters were then isolated as *Pac*I fragments from their respective vectors, and subsequently cloned into binary vector PIPRA560 [35].

**Figure 1:**
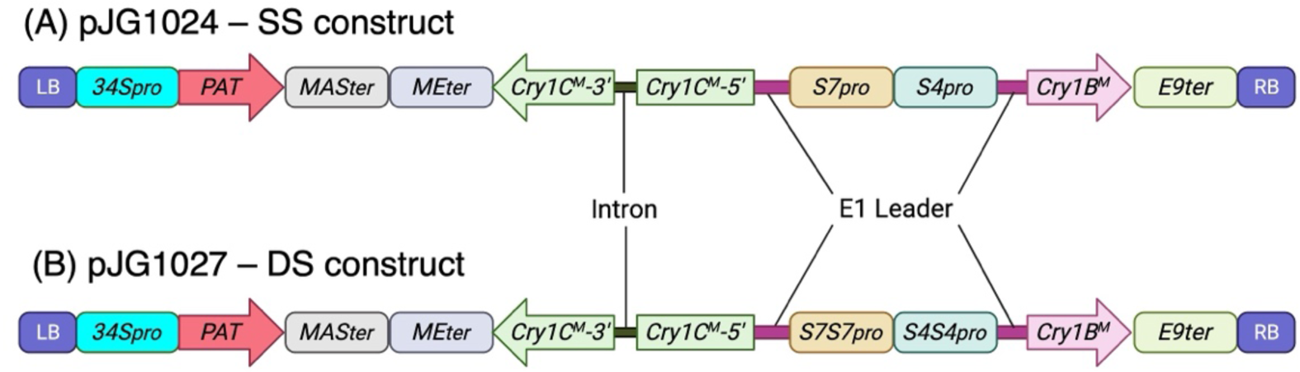
Schematic diagram of T-DNA region of constructs used to test the insecticidal activity of *Cry1B*^*M*^ and *Cry1C*^*M*^ genes in plants; *34S*_*pro*_: Promoter of Figwort mosaic virus (FMV) *34S* RNA gene [36]; *MAS*_ter_: *Agrobacterium tumefaciens MANNOPINE SYNTHASE* (*MAS*) gene terminator [35]; *Cry1B*^*M*^/*Cry1C*^*M*^: Modified *Cry1B*/*Cry1C* genes; *S4*/*S7*: Subterranean Clover Stunt Virus *S4* and *S7* promoters [34]; *E9*_*ter*_: Terminator region of the pea *Rubisco E9* gene [35]; *ME*_*ter*_: Terminator region of *Flaveria bidentis MALIC ENZYME* (*ME*) gene [34]; Intron: second intron (IV2) of the potato gene *ST-LS1* [36], E1: The 5’ leader sequence of the tapetum specific *E1* gene of *Oryza sativa*, LB: Left border; RB: Right border.

A *NPTII* expression cassette comprising a figwort mosaic virus *34S* promoter [37], the coding sequence of the *NEOMYCIN PHOSPHOTRANSFERASE II* (*NPTII*) gene [38] and the terminator of *Agrobacterium MANNOPINE SYNTHASE* (*MAS*) gene (*MAS*_*ter*_) [35] was synthesized and cloned into the *Eco*RI/*Hin*dIII sites of the pUC19 vector. A glufosinate-ammonium resistant selectable marker was generated by replacing *NPTII* with the *PHOSPHINOTHRICIN ACETYLTRANSFERASE* (*PAT*) gene [39]. The PAT selection cassette was then isolated from this plasmid and cloned into the *Sac*II site of the binary vectors containing the modified *Cry* genes under the control of either *S7-S4* or *S7S7-S4S4* stunt promoters. These constructs, pJG1024 (single stunt (SS) construct) and pJG1027 (double stunt (DS) construct) (Figure 1) were then introduced into *Agrobacterium* (C58) via electroporation.

### Plant Transformation

pJG1024 and pJG1027 were inserted into wild-type *Arabidopsis thaliana* using floral dipping [40]. Transgenic plants were identified using glufosinate-ammonium (100 μg/ml) selection on soil and further confirmed by amplifying the herbicide resistance gene *PAT* using primers BaR-F (5′-GTTGGTTGCTGAGGTTGAG-3’) and BaR-3’
sR (5′-TGGGTAACTGGCCTAACTG G-3′). For each construct, ten independently transformed T_1_ lines were randomly selected, and their progeny exposed to glufosinate-ammonium selection. Based on segregation ratios, lines judged to have a single T-DNA insertion (1:3, Glufosinate ammonium sensitive: Glufosinate ammonium resistant) were selected for further analysis. Homozygous T_2_ plants were subsequently isolated for each of the T_1_ deemed to have a single segregating T-DNA insertion and were used in all subsequent assays.

### Insect Bioassay

A colony of diamondback moth (DBM) susceptible to the Bt toxins encoded by *Cry1*^*M*^ genes were maintained in an insect growth chamber at 25°C. Arabidopsis plants homozygous for the T-DNA insertion were grown for ∼ four weeks and their mature leaves collected for insect bioassay, RNA extractions, and protein quantification. For insect bioassay, two leaves were placed on a moist filter paper in a plastic cup (size 40 × 50 mm). On each leaf, ten DBM larvae (1^st^ instar) were deposited. Larval mortality and leaf damage were scored after 48 hours and at 72 hours in cases where the larvae survived after 48 hours. Insect bioassays were performed at 25°C. Leaf damage was scored on scale from 0 (no visible damage) to 5 (leaf skeletonised).

### Quantification of Cry1C^M^ protein

The abundance of Cry1C^M^ protein in leaves of transgenic plants was quantified using a Cry1C-specific enzyme-linked immunosorbent assay (ELISA) assay (Cry1C-specific Quantiplate Kit; Envirologix, USA). Briefly, leaves collected from transgenic lines were weighed and put into a 1.5 ml tube. The tubes were then placed in Ziplock plastic bags containing silica beads and dried over a period of two weeks. Proteins were extracted from 1 mg dried tissue using the extraction buffer supplied with the kit. The ELISA was performed according to the manufacture’s instruction. The amount of expressed Cry1C^M^ protein in the leaf sample was calculated from a standard curve generated using the pure Cry1C protein supplied with the Quantiplate ELISA kit. The amount of Cry1C^M^ protein content in the samples was determined using the standard curve and given as ng per gram fresh weight (FW) of leaves.

### RNA extraction and reverse transcription quantitative PCR (RT-qPCR)

RNA was extracted from leaf tissue using a Spectrum Plant Total RNA kit (Sigma, USA) according to the manufacturer’s protocol. Extracted RNA was treated using Turbo DNA-free kit (Ambion, USA) to remove contaminating genomic DNA before first strand cDNA was generated using Oligo (dT) primers and Superscript III reverse transcriptase (Thermo Fisher). RT-qPCR was performed using a SensiMix SYBR No-ROX Kit (Meridian, Australia). Briefly, 10 μl qPCR reactions containing 1μl diluted (1:10) cDNA, 2.5 μM forward and reverse primer, with 1x SYBR Green Master Mix were set up in triplicate and run on a Bio-Rad CFX96 real time PCR machine. Cycle threshold (Ct) values were calculated using the Bio-Rad CFX manager version 3.1. The relative *Cry1B*^*M*^*/Cry1C*^*M*^ mRNA expression level were determined using the comparative Ct method and normalized to *ACTIN2* (*AT3G18780*). The sequences of RT-qPCR primer used in this study were, Actin2-F (5′-TCTTCCGCTCTTTCTTTCCA-3′), Actin2-R (5′-TCTTCCGCTCTTTCTTTCCA-3′), Cry1B-F (5′-TAGAGGGACCGCTAACTATT C-3′)/Cry1B-R (5′-CGACAACCGATGTGAGTAAG-3′), and Cry1C-F (5′-GAAAGAATGCCGCAA TGTC-3′)/ Cry1C-R (5′-CTTACAACCGTGGGCTTAAC-3′).

### IP landscape analysis

IP searches were performed using keywords and sequence-based approaches to identify relevant patent filings in national databases for filings in the US (https://ppubs.uspto.gov/pubwebapp/static/pages/ppubsbasic.html), Australia (https://www.ipaustralia.gov.au/), EPO (https://www.epo.org/en/searching-for-patents/technical/espacenet) and WIPO (https://patentscope.wipo.int/search/en/search.jsf), and India (https://iprsearch.ipindia.gov.in/publicsearch). This provided information about the legal status of patents as well as their file histories. Results from these searches are provided in Table 2 and Supplementary Table 1.

**Table 1.**
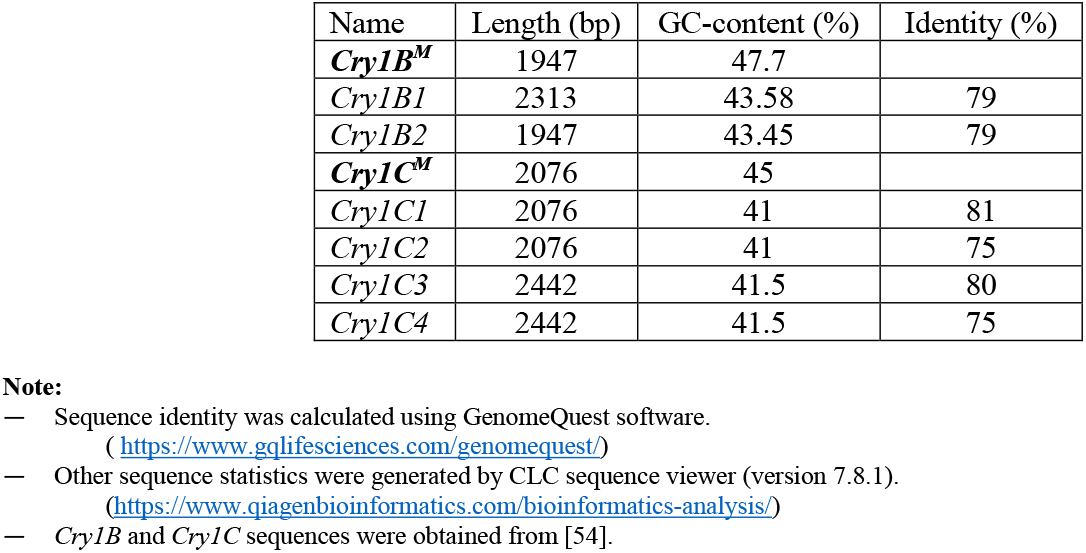
GC-content of *Cry1B*^*M*^/*Cry1C*^*M*^ nucleotide sequences compared to other *Cry1B*/*Cry1C* nucleotide sequences.

**Table 2.**
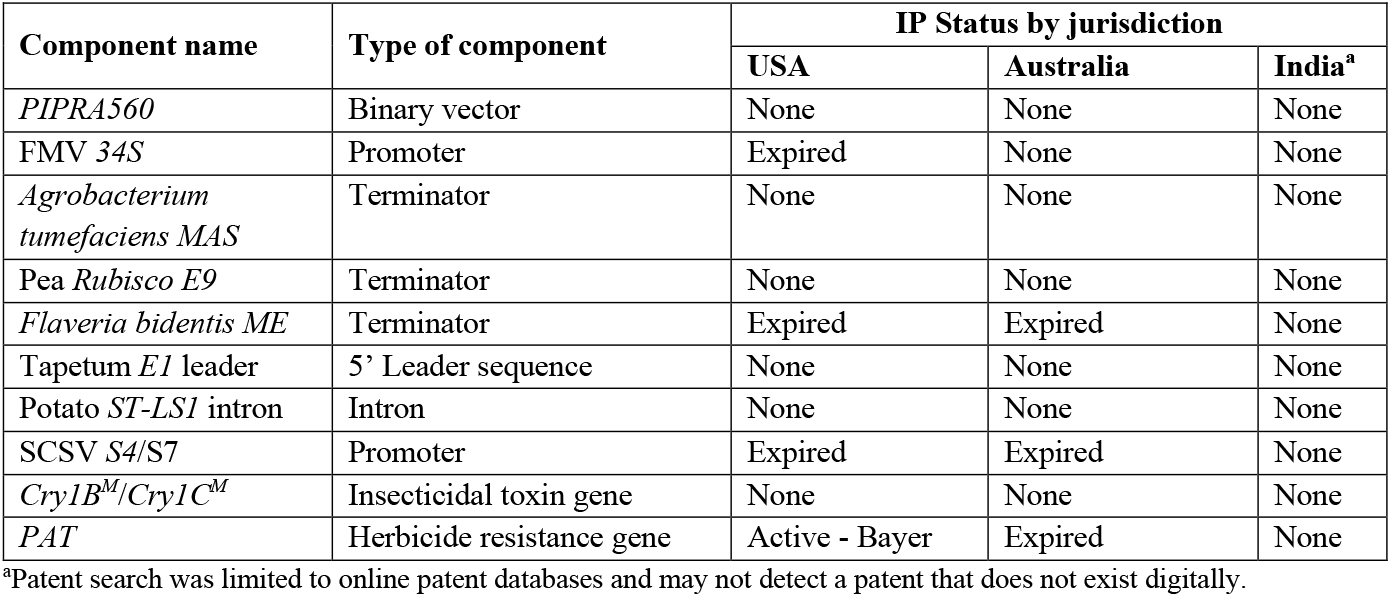
IP status of the components selected for the <u>Bt</u> gene constructs described in this study.

## 3.0 Result

### Modification of *Cry1B* and *Cry1C* genes used in this study

To maximize activity of *Cry1B*/*Cry1C* genes in plants, we modified the *Cry1* sequences (*Cry1*^*M*^) to eliminated features that are known to reduce the expression of these genes in eukaryotic cells. This included extensive codon-optimization, which involved the selection of codons used at high frequently in *Brassica* genes and are GC-rich [41]. Following this, the predicted GC-content of the modified *Cry* genes *Cry1B*^*M*^ and *Cry1C*^*M*^ was 47.7% and 45%, respectively, which is higher than in their unmodified versions (Table 1).

All alternative reading frames (ORFs) were removed from the modified *Cry* genes, a key requirement for GM plants needing regulatory approval before commercialisation. Finally, all sequences that are known to affect transcript stability, such as ATTTA and splice donor and acceptor site such as AG and GT, respectively, were removed from the modified *Cry* gene sequences along with any internal polyadenylation signal sequences that might cause premature termination of transcription. Following these modifications, the degree of identify between known *Cry1B* genes (e.g., *Cry1B1* and *Cry1B2)* and *Cry1B*^*M*^ was 79% at DNA level (Table 1) and the identity between *Cry1C*^*M*^ and four other *Cry1C* genes ranged from 75-81% (Table 1).

### Design of *Cry1B*^*M*^/*Cry1C*^*M*^ constructs

Where possible components and methodologies that are free of third-party IP were used in the development of the *Cry1*^*M*^ constructs to minimize IP obstacles, including any licensing costs associated with eventual commercial cultivation of plants expressing the *Cry1*^*M*^ genes. We performed a detailed online database search of the patents surrounding the binary vector, promoters, terminators, selectable markers, *Cry* genes and methodologies used in the generation of the Bt constructs and list their current IP status in Australia, USA, and India in Table 2 and Supplementary Table 1.

The *Cry1B*^*M*^/*Cry1C*^*M*^ gene construct was designed so that physical linkage between the genes ensured that they integrate into the same chromosomal site following transformation (Figure 1). This design eliminated the need for crossing to combine transgenes following their separate introduction into plants. Binary vector PIPRA560 was selected for use in these experiments as it is available under royalty-free terms for commercial cultivation in developing countries, including India and under modest fee-based terms for developed countries [35]. The herbicide resistance gene *PAT* was chosen as a plant selectable marker for transgenic plant selection as it had FTO in both Australia and India. The FMV *34S* promoter [37] and the terminator region of the *MAS* [35] gene were placed upstream and downstream of the *PAT* gene, respectively. These components are present within the PIPRA560 plasmid and were obtained under a UC Davis licensing agreement. Subterranean clover stunt virus (SCSV) promoters *S4* and S7 were selected because previous work had shown that their use with other *Cry* genes led to high-level expression and subsequent insecticidal activity [34]. These are now available free of third-party IP (see Table 2). Two different configurations of these promoters were tested; the first being single *S4*/S7 promoters (Figure 1A; hereafter referred to as SS) and the second being a double promoter configuration (Figure 1B; hereafter referred to as DS). By analysing expression of the *Cry1*^*M*^ genes arising from SS and DS constructs, we addressed whether these promoters arranged in tandem conferred a significantly higher level of expression than a single promoter configuration, as suggested in previous studies of these promoters [34]

### Generation of transgenic lines and insect bioassay

More than 40 independent T_1_ plants transformed with a T-DNA containing either the SS cassette or a DS cassette were generated. Of these ten SS and DS primary transformants were randomly selected for initial insect bioassays and Cry1C protein content analysis (data not shown).

First instar DBM larvae were placed on leaves collected from these primary transgenic Arabidopsis lines hemizygous for the T-DNA, along with those from wild-type plants. Larvae fed on the wild-type leaves developed normally, resulting in severe leaf damage associated with unconstrained feeding (Figure 2A). In contrast, transgenic leaves remained undamaged from larval feeding (Figure 2A). The number of live and dead larvae were assessed (Table 3).

**Table 3.** Insecticidal mortality and degree of feeding damage seen on leaves derived from wild-type and primary (T_1_) transgenic Arabidopsis lines hemizygous for the *Cry1B*^*M*^/*Cry1C*^*M*^ construct.

| Mortality of larvae (%) | Single stunt (n=10 independent lines) | Double stunt (n=10 independent lines) | Untransformed control (n=2) |
| --- | --- | --- | --- |
| 24 hrs | 98.5 | 99.5 | 0.00 |
| 48 hrs | 100 | 100 | 0.00 |
| 6 days | 100 | 100 | 31.5 |
| Plants with live larvae (%) |  |  |  |
| 24 hrs | 15 | 4 | 100 |
| 48 hrs | 0 | 0 | 100 |
| 6 days | 0 | 0 | 100 |
| Live larvae moulted beyond 1<sup>st</sup> instar at 4 days | - | - | >40% |
| Mean leaf damage score at 4 days<sup>a</sup> | 0.125 | 0.125 | 4.2 |

<sup>a</sup>Leaf damage score was assessed as below:
- 0 – No visible sign of damage - 1 – Slight scrapping of leaf surface - 2 – Small holes through leaf - 3 – Large holes through leaf - 4 – Widespread leaf damage - 5 – Skeletonised

**Figure 2:**
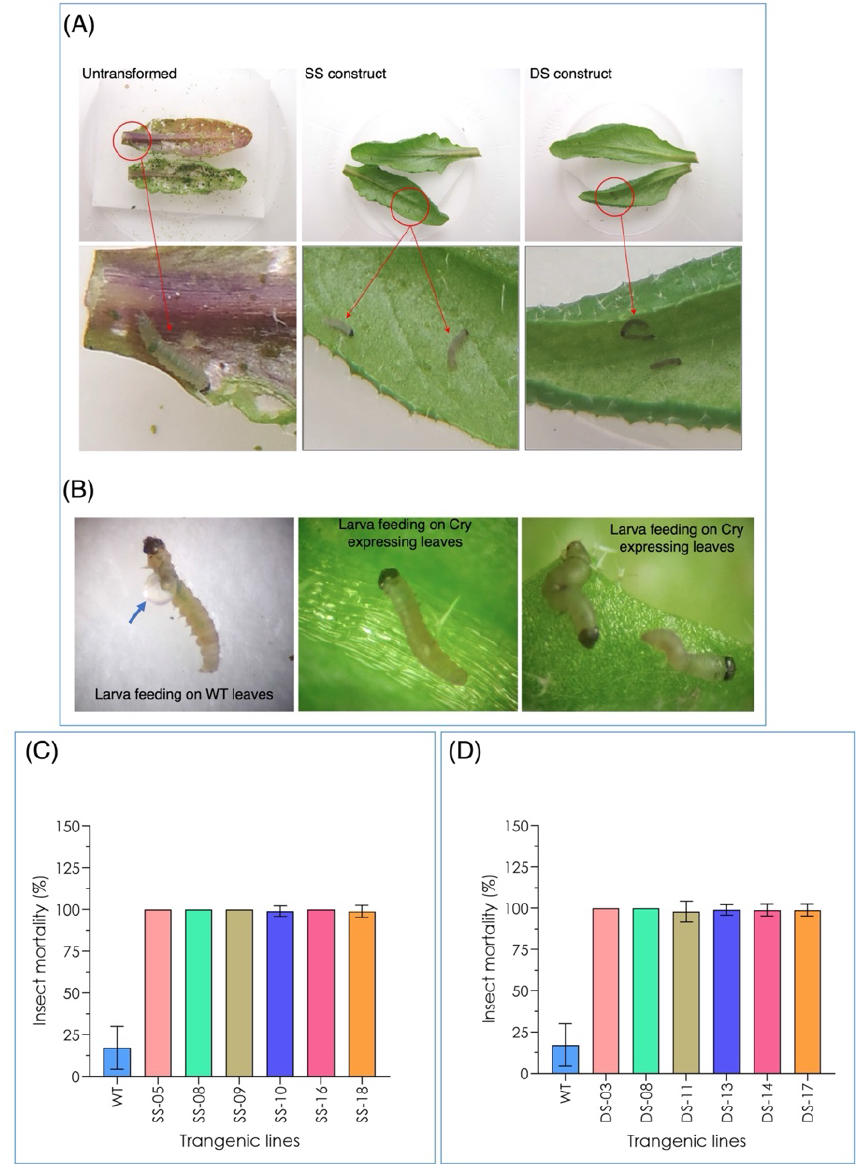
Insect bioassay on Arabidopsis leaves derived from transgenic plants homozygous for *Cry1B*^*M*^/*Cry1C*^*M*^ transgene. (A) Transgenic leaves expressing *Cry1B*^*M*^*/Cry1C*^*M*^ genes under a single stunt (SS) and double stunt (DS) *S4* and *S7* promoters. (B) Image showing insect larval guts (indicated with blue arrowhead) after feeding on wild-type leaves. (C) Insect mortality found in individual transgenic lines having *Cry1B*^*M*^/*Cry1C*^*M*^ expression under a single stunt (*S4*/*S7*) promoter. (D) Insect mortality associated with in individual transgenic lines with *Cry1B*^*M*^/*Cry1C*^*M*^ expression under the control of double stunt (*S4S4*/*S7S7*) promoters.

After 24 hours, approx. 99% larvae placed on the T_1_ leaves were dead, whereas all larvae placed on the wild-type leaves were alive and actively feeding (Table 3). After 48 hours, all remaining larvae feeding on the transgenic leaves were dead, while almost all the larvae feeding on the wildtype leaves were alive (Table 3). Significant larval death (31.5%) was seen on the wildtype leaves, but only after day 6. Moreover, ∼40% larvae placed on the wild-type leaves were found to have moulted beyond 1^st^ instar, which was not observed for larvae placed on transgenic leaves. During the insect bioassay, the health of the larvae was examined. Healthy larvae were present on the wild-type leaves, whereas those feeding on the transgenic leaves appeared shrivelled and small, including some displaying gut bursting (Figure 2B). Segregation analysis performed on each of these ten primary transformants identified six with a single segregating T-DNA insertion. Homozygous T_3_ progeny derived from these lines were subsequently used for insect feeding assays, which revealed 100% mortality within 48 hours of feeding on transgenic leaves (Figure 2C, D). Interestingly, there was no discernible difference in insect mortality between transgenic leaves expressing *Cry1*^*M*^ genes under a single stunt promoter from those under a double stunt promoter (Figure 2C, D).

### *Cry1B*^*M*^ *and Cry1C*^*M*^ expression level in the plants

Expression of *Cry1B*^*M*^ and *Cry1C*^*M*^ in vegetative tissue of seedlings homozygous for the SS and DS constructs was measured by qRT-PCR. While this revealed expression of both transgenes in all plants (Figure 3A, B), considerable variation was observed. For instance, lines SS-08 and DS-08, displayed high levels of both *Cry1B*^*M*^ and *Cry1C*^*M*^ expression, whereas low expression of both genes was detected in lines DS-13 and DS-14 (Figure 3A, B). Except for SS-08, SS-09 and DS-08, DS-17, most transgenic lines (n=12) displayed significant differences between the *Cry1B*^*M*^ and *Cry1C*^*M*^ expression with the majority having higher *Cry1B*^*M*^ expression compared to *Cry1C*^*M*^ (Figure 3A, B).

**Figure 3:**
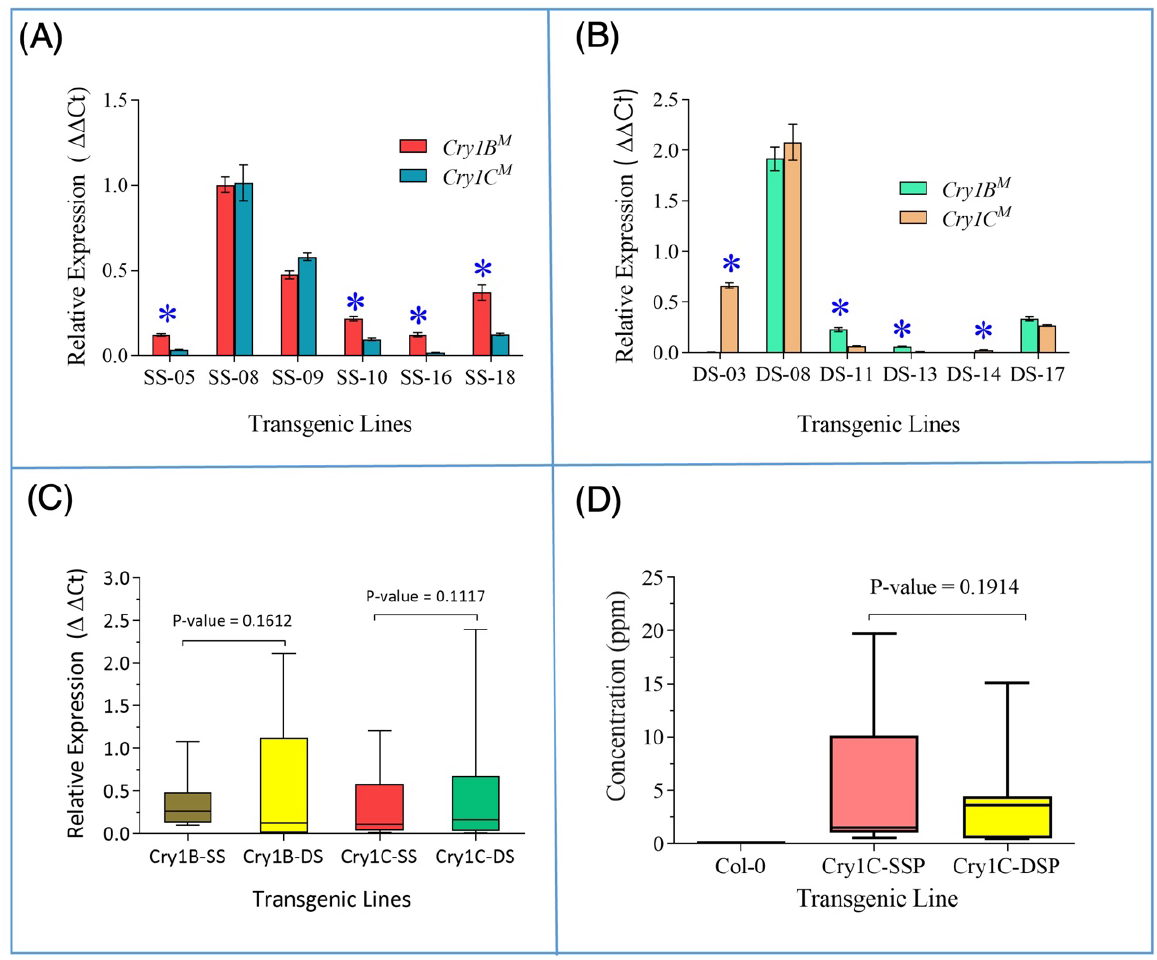
Expression of *Cry1B*^*M*^ and *Cry1C*^*M*^ transgenes in the transgenic Arabidopsis plants. (A) *Cry1B*^*M*^*/Cry1C*^*M*^ expression seen in transgenic lines with a single stunt (*S4/S7*) promoter (six independent transgenic events examined with 5 plants per event). (B) *Cry1B*^*M*^*/Cry1C*^*M*^ expression in transgenic lines with double stunt (*S4S4/S7S7*) promoters (six independent transgenic events examined with 5 plants per event). (C) Comparison of *Cry1B*^*M*^ and *Cry1C*^*M*^ expression (average of six independent transgenic events) under *S4/S7* single and double stunt promoters. A paired *t*-test found no statistical differences in the range of *Cry* gene expression values seen in plants with single or double stunt promoters. (D) Comparison of Cry1C^M^ protein content in plants with single or double stunt promoters as assessed by enzyme-linked immunoabsorbent assay (ELISA) (n=3 plants). SS: Lines with single stunt promoters, DS: Lines with double stunt promoters. Lines denoted by “*” indicates expression of *Cry1B*^*M*^ and *Cry1C*^*M*^ is significantly different. Statistical significance was assessed by multiple *t* test based on the Holm-Sidak method [53].

Levels of Cry protein was quantified by ELISA (Table 4). This analysis was restricted to Cry1C^M^ due to the unavailability of a Cry1B^M^-specific ELISA kit. For SS lines, the quantity of Cry1C^M^ protein ranged from 0.81 to 17.69 μg/g leaf fresh weight (FW) with significant differences in protein content between transgenic lines (F = 87.26, p < 0.0001).

**Table 4.** Cry1C^M^ protein content of individual T_3_ transgenic lines homozygous for the T-DNA that were used for insect bioassay.

| *Cry1B<sup>M</sup>/Cry1C<sup>M</sup>*<br>expressing lines | $\mu\text{g Cry1C}^{\text{M}}$ protein/g leaf FW<br>(mean ± SEM), (n = 3 plants) |
| --- | --- |
| Wild type (Col-0) | 0.01 ± 0.002 |
| SS-05 | 1.64 ± 0.097 |
| SS-08 | 17.69 ± 1.574 |
| SS-09 | 9.95 ± 0.780 |
| SS-10 | 1.53 ± 0.383 |
| SS-16 | 0.81 ± 0.133 |
| SS-18 | 0.99 ± 0.238 |
| DS-03 | 3.50 ± 0.489 |
| DS-08 | 13.48 ± 0.822 |
| DS-11 | 4.01 ± 0.680 |
| DS-13 | 0.51 ± 0.027 |
| DS-14 | 0.52 ± 0.018 |
| DS-17 | 3.77 ± 0.158 |

SS: *Cyr1B<sup>M</sup>/Cry1C<sup>M</sup>* under single stunt promoter DS: *Cyr1B<sup>M</sup>/Cry1C<sup>M</sup>* under double stunt promoter FW: Fresh weight

Similarly, Cry1C^M^ protein in DD lines ranged from 0.51 to 13.48 μg/g leaf FW with significant differences also detected between lines (F = 97.29, p < 0.0001). It is worth noting that a previous study using a leaf-dip assay with pure Cry1Ca4 protein found that the lethal concentration (LC_50_) to be <1.18 ppm when fed to 26 global DBM populations and an average of only 0.18ppm in Indian DBM populations [42]. This suggests that most of transgenic lines generated in this study had Cry1C protein levels that would by themselves be effective against DBM (Figure 3D).

Initial observations failed to detect a noticeable difference in the range of insecticidal activity displayed by transgenic lines expressing *Cry1B*^*M*^/*Cry1C*^*M*^ under the control of single or double stunt promoters (Figure 2C, D). Consistent with this observation, no significant differences were found in the expression of lines transformed with SS as opposed to DS constructs as measured by qRT-PCR of the modified *Cry* genes (*Cry1B*^*M*^, p-value 0.161; *Cry1C*^*M*^, p-value 0.112). Similarly, the range in Cry1C^M^ protein content in leaves of SS and DS lines did not differ significantly (p-value 0.191).

### Correlation between the *Cry1C*^*M*^ expression and protein content

To determine the relationship between the amount of *Cry1C*^*M*^ mRNA and corresponding protein content, qRT-PCR and ELISA results were compared. In most cases, levels of mRNA corresponded closely to protein content (SS-08, SS-09, DS-14, DS-17; Figure 4A). Furthermore, a Spearman rank correlation coefficient test identified a strong statistical correlation between the Cry1C^M^ protein content and *Cry1C*^*M*^ transgene expression (Rs = 0.0846, P = 0.001). While there was a good correlation between gene expression and protein levels, there were some notable exceptions. For instance, the highest amount of *Cry1C*^*M*^ transcript was detected in line DS-08 but this did not correlate with the highest amount of detectable Cry1C protein. Conversely, the high level of Cry1C protein content in line SS-08 arose from transcript levels of *Cry1C*^*M*^ that were nearly half of that observed in DS-08 (Figure 4A).

**Figure 4:**
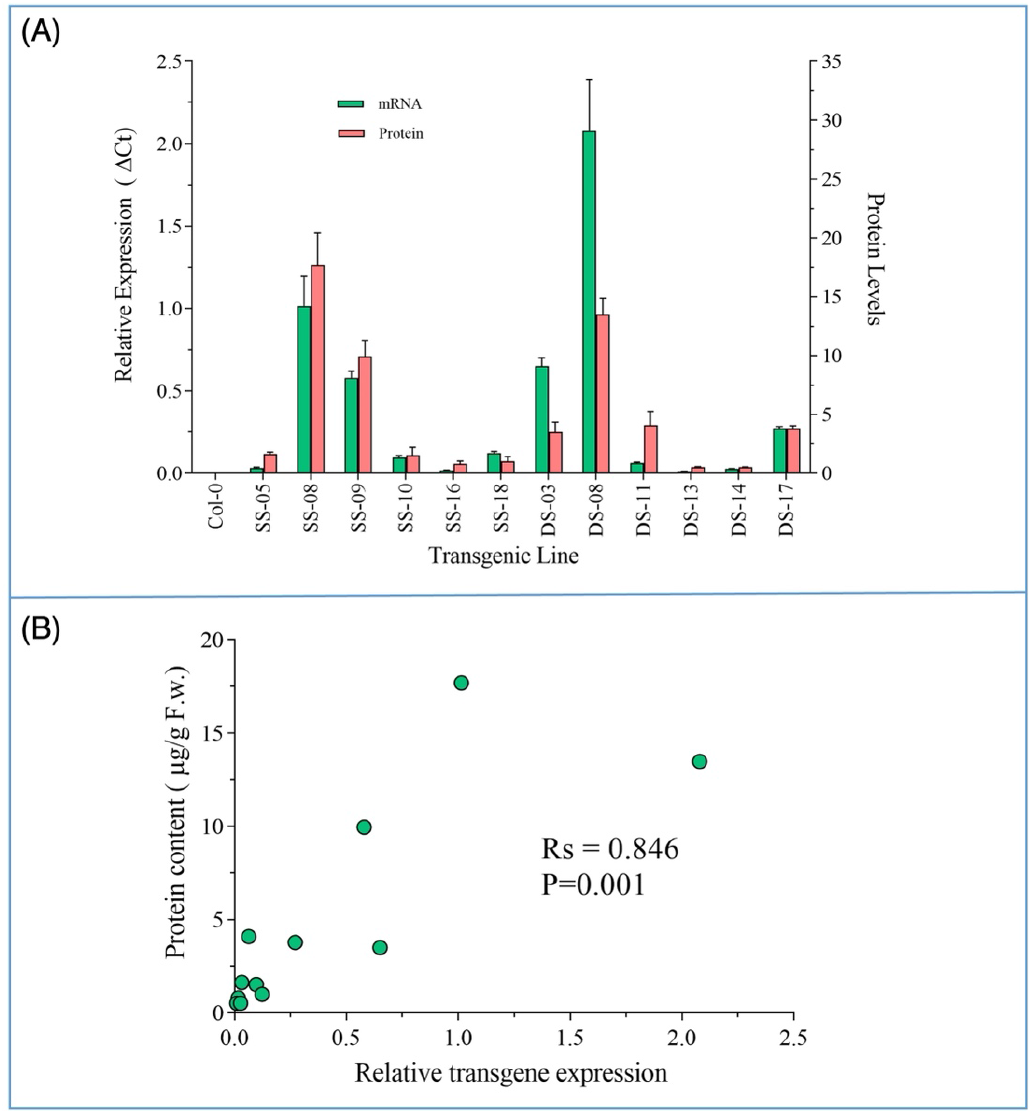
Comparison of *Cry1C*^*M*^ transgene expression and protein accumulation in transgenic Arabidopsis lines. (A) Histogram showing the amount of *Cry1C*^*M*^ mRNA and protein content in twelve transgenic lines tested. (B) Spearman correlation between relative transgene expression and Cry1C^M^ protein content in the lines listed in Figure 2 C and D. SS: Lines with single stunt *S4/S7* promoters; DS: Lines with double stunt *S4S4*/*S7S7* promoters.

## 4.0 Discussion

The *Cry1B*/*Cry1C* combination, whose modification is described here, was previously shown to be effective against DBM and has never been used in commercially available GM crops or as the basis for sprayable Bt insecticides [43, 44]. For instance, purified Cry1Ba2 and Cry1Ca4 proteins display LC_50_ values <0.91 ppm and <1.18 ppm, respectively, when tested against of DBM populations [42]. Furthermore, no cross-resistance was found between Cry1Ba2 and Cry1Ca4, or in experiments aimed at generating resistance to the two Bts in DBM populations [43]. The minor resistances that were observed in these studies were unstable and genetically recessive, as well as being associated with a high fitness costs [43]. Where resistance to Cry1C in DBM has been identified, it manifests as a polygenic trait [45]. Taking these observation together with the fact that Cry1B and Cry1C bind to different receptors in the insect gut [44], suggests that insect resistance to this combination of *Cry* genes is unlikely to arise.

While previous work [43, 44] found that transgenic *Brassica* expressing the *Cry1B*/*Cry1C* stack displayed robust resistance to a range of lepidopteran pest species in small trials run in India, these lines (developed by a public/private-funded consortium), were not developed further due to the length of time required to gain regulatory approval for commercial planting. Despite this set-back, the Australian and Indian public partners in this consortium wanted to continue the development of Brassica expressing Bt toxins for their respective markets. To facilitate this, it was necessary to alter the sequence of the *Cry* genes so that they were unequivocally not that of the private partner. We used this opportunity to both free the new *Cry1*^*M*^ gene constructs of proprietary IP as far as practicable and to optimise the nucleotide structure of the *Cry* genes for expression in plants. The PIPRA560 plant binary vector was used to deliver the *Cry* constructs to plants. The patent holder, the University of California, Davis, allowed licence free research use and free commercialisation rights for developing countries, including India. The terms of use state that any construct developed using PIPRA560 must also be free to use by others.

As stated previously, one way to overcome the development of resistance to Bt toxin is to express the *Cry* gene at high level so that insects heterozygous for resistance mutation are eliminated from the population. Given this, we choose double stunt SCSV promoters for our Bt toxin gene construct design as plants expressing *Cry* gene under two stunt promoters would be more effective in killing insect than their single stunt counterpart. Interestingly, the results reported here show that there is no obvious difference in insect mortality between the plants expressing *Cry1*^*M*^ genes under single or double stunt SCSV promoters (Figure 2C, D; Table 3). Furthermore, previous work characterising double stunt promoters has indicated that *S7S7* double promoter was better than *S4S4* promoter, whereas our results indicated that *S4S4* is slightly more active than the *S7S7* promoter (Figure 3C). Differences between our study and those previously might reflect the assay system used to compare the promoter strength. For instance, the earlier studies of the *S4S4* and *S7S7* promoters relied upon *Cry1Ab* protein content assays to measure activity of the promoters, whereas in our study we used a combination of both protein and mRNA assays. As mentioned above protein production from mRNA is affected by multiple factors and hence protein content in GM plants may not be a reliable indicator of promoter strength. As performance of *S4S4* and *S7S7* double stunt promoters also varied according to plant species used for transformation (cotton, tobacco and tomato; [34]), variation in SCSV promoter activity observed in our study might also reflect background differences between Arabidopsis and plants used in earlier studies.

Despite variation in *Cry1B*^*M*^ and *Cry1C*^*M*^ expression between and within lines, there was no detectable variation in insecticidal activity, as 100% insect mortality was achieved within the 48 hours even in lines with low *Cry1B*^*M*^*/Cry1C*^*M*^ expression (Figure 1B, C). Although transcript levels were measured both for *Cry1B*^*M*^ and *Cry1C*^*M*^, quantification of Cry1B protein could not be performed due to lack of a Cry1B*-*specific ELISA commercial kit. Unfortunately, lack of access to the original *Cry* gene constructs prevented us from directly testing whether the modified sequences represent a significant improvement over the insecticidal activity of the unmodified sequences. Despite this, it seems reasonable to conclude that both proteins retained insecticidal activity under lab conditions. This is inferred from the observation that in some lines, one *Cry* gene was expressed at much higher levels than the other (e.g., SS-05, SS-16 DS-03; Figure 3B), yet still conferred 100% DBM larvae mortality. While there was a clear correlation between *Cry1C*^*M*^ gene expression and protein abundance (Figure 4), there were a few notable exceptions, e.g., line DS-03 and DS-08. This discrepancy presumably reflects inefficient conversion of mRNA to protein, an observation that has been reported in several other studies of *Cry* transgene activity [46, 47]. However, due to a relatively modest sample size (n = 12 lines) in our study, as well as those reported in others [46, 47], it is difficult to discern a clear relationship between *Cry* gene expression and protein accumulation.

A potential problem arising from gene stacks is that when more than one gene is placed in the same T-DNA, their expression may be compromised due to gene silencing, particularly if they share similar sequences and regulatory elements such as promoters, 5’ s-UTRs and 3′-UTRs [48]. Therefore, expression analysis of each gene in plants is important, as silencing or sub-optimal expression of one gene may result in reduced efficacy of the insecticidal protection provided by the *Cry* gene stack. We found substantial differences in expression levels of *Cry1B*^*M*^ and *Cry1C*^*M*^ in most of the lines (Figure 3A, B). The variation in *Cry1B*^*M*^/*Cry1C*^*M*^ expression seen in the same line might reflect issues associated with having the stunt promoters directly adjacent to one another in reverse orientation within the T-DNA. Such an arrangement may make these transgenes prone to gene silencing, which would reduce their efficacy. Alternatively, differences in *Cry* gene expression might reflect position effects, in which genomic regions adjacent to the T-DNA insertion site influence transcription activity of the transgenes. Similarly, variation in the extent of T-DNA insertion or its rearrangement prior to or after integration into the genome may influence *Cry* gene activity. Such variation between lines transformed with the same construct has been previously reported [49] and thus is not without precedent. Importantly, variations in *Cry1B*^*M*^ and *Cry1C*^*M*^ expression level observed in the transgenic Arabidopsis lines do not seemingly reflect an issue with their coding sequences. This can be inferred from the fact that in some cases *Cry1B*^*M*^ is expressed at higher levels than *Cry1C*^*M*^ (e.g., SS-18, DS-11), and vice versa (e.g., SS-09, DS-03).

The results provided here illustrate the types of sequence modification that can be successfully introduced into *Cry* genes as well as the suitability of components chosen for constructs that have FTO such as the PIPRA560 vector. Other than the *Cry* genes themselves, the components used in the gene constructs reported here are to the best of our knowledge currently free of third-party IP in Australia and India. Confirmation that this applies in other countries would require detailed patent searches to be undertaken. The use of components that have FTO in both the research and the commercialisation phases in public GM breeding programs is important as it can substantially reduce the complexities and costs faced in the commercialisation phase [35]. While the work here only reports the activity of the *Cry1B*^*M*^/*Cry1C*^*M*^ gene stack in the model plant Arabidopsis under laboratory conditions, work was undertaken to introduce these constructs into elite Brassica crop lines, and preliminary analysis suggests that they are as effective in crop plants as they are in Arabidopsis [50]. Unfortunately, funding constrains prevented these transgenic crop lines from being fully assessed for insect resistance in field trials.

## 5.0 Conclusion

Despite the obvious benefits of transgenic plants expressing *Cry* genes, which include preventing large scale crop losses from insect attack, this technology has been applied to only a few crops such as cotton, canola and maize which are grown on a large enough scale to make the costs of deregulation and the separation of the product in harvesting, storing and marketing economically attractive [51]. The timelines, costs, and political opposition to GM crops in a significant proportion of markets has delayed the introduction of *Cry* transgenes into other crops such as vegetables and major grain crops such as rice or wheat. Furthermore uptake of Bt technology in developing nations has been severely curtailed by the difficulties of access to IP held by entities such as multinational seed companies [52]. Given this, publicly funded research organizations and academic institutions in developing nations have an incentive to develop their own Bt crops [35] in which licencing issues associated with the use of genetically modified material is kept low so that farmers can take advantage of the considerable benefits arising from the technology [51]. Our work here provides an example of an approach that might be taken to achieve this aim.

## Material

The PIPRA560 vector was obtained from PIPRA, Department of Plant Sciences, University of California, Davis, CA95616.

## Author contributions

MMH, DAR, CR and JG conceived the idea. MMH and AW undertook the molecular biology and plant work; FT and DAR conducted the insect experiments. JC and DAR conducted the IP analysis. MMH led the writing and revision of the manuscript. All authors accepted the final version of the manuscript.

