## Supplementary Table 1 for "Designing Bt constructs for Brassicas, with minimal IP issues – A case study"

| | Technology inventory | Use of technology | IP status by jurisdiction | | | IP information | Relevant patents/publications |
|---|---|---|---|---|---|---|---|
| | | | USA | Australia | India | | |
| **I. Plant Transformation DNA Plasmid Backbone** | | | | | | | |
| **Binary Vector** | pPIPRA560 | Backbone Plasmid | **None** | **None** | **None** | Use of PIPRA560 does not fall under a patent. However as DNA was obtained from UC Davis, its use is covered by a MTA that was executed in Nov 2012 with the University of Melbourne, which limits use of the plasmid for academic non-commercial research.<br><br>Note: Entities wanting to use the PIPRA560 vector for humanitarian use can negiote with UC Davis, as this institution provides access to these materials under royalty-free terms for humanitarian use and under fee-based terms for commercial use. | Chi-Ham, C. L. et al. (2012). "An intellectual property sharing initiative in agricultural biotechnology: development of broadly accessible technologies for plant transformation." Plant Biotechnol J 10(5): 501-510. |
| **II. T-DNA Region** | | | | | | | |
| **Plant selectable marker cassette** | FMV *34S* | Promoter | **Expired** | **None** | **None** | Use of the FMV *34S* promoter falls under the patent "Figwort mosaic virus promoter and uses" assigned to Calgene Inc. (now Monsanto) and the University of California. UC Davis solely manages the Intelectual property (IP) and Tangible property (TP) rights. The US patent expired April 18th, 2017. The US patent family does not apparently contain members in Australia or India. | US6051753 |
| | MAS 3'UTR | Transcription Regulation, 3'UTR | **None** | **None** | **None** | No relevant IP identified. | |
| **BT *Cry1C* DNA Cassette** | SCSV *S7* or *S7S7* | Promoter | **Expired** | **Expired** | **None** | Use of the SCSV *S4* and *S7* promoters fall under two patent families: "Plant transcription regulators from circovirus" assigned to CSIRO and "Novel Genes Encoding Insecticidal Proteins" assigned to BASF Agricultural Solutions Seed US LLC (formerly Bayer Bioscience). The Australian CSIRO patent AU689311 and US6211431 expired August 30th, 2015. The BASF Agricultural Solutions Seed US LLC patents expire 16th march 2027.<br><br>The BASF patent family covers the specific use of the subterranean clover stunt virus SCSV *S7* or *S4* promoters to drive the *Cry1C* or *Cry1B* gene expression. Due to a narrowing of claims during prosecution of the patent in both the US and Australia (see *Cry1B* /*Cry1C* entry below), it is likely that use of the SCSV promoters in conjunction with *Cry1*$^{M}$ sequences falls outside of these claims.<br><br>No IP was found in the Indian online patent databases. | CSIRO: US6211431, AU689311<br>Bayer: AR059995; AT511515; AU2007228981; CA2646471; CN101405296; DK1999141; EA200802018; EP1999141; NZ571952; US20100235951; WO2007107302 |
| | *Oryza sativa* tapetum *E1* (GE1) | Transcription Regulation, 5' UTR leader | **None** | **None** | **None** | Use of tapetum *E1* (GE1) Oryza sativa 5' UTR leader sequence falls under the patent family "Novel Genes Encoding Insecticidal Proteins" assigned to BASF Agricultural Solutions Seed US LLC. Due to a narrowing of claims during prosecution of the patent in both the US and Australia (see Cry1B/Cry1C entry below), it is likely that use of *E1* in conjunction with *Cry1*$^{M}$ sequences falls outside of these claims. AU2007228981, EP1999141 and WO2007107302 applications were filed March 16th, 2007 and will expire March 16th 2027.<br><br>No IP was found in the Indian online patent databases. | AR059995; AT511515; AU2007228981; CA2646471; CN101405296; DK1999141; EA200802018; EP1999141; NZ571952; US20100235951; WO2007107302 |
| | Potato intron | Transcription Regulation intron | **None** | **None** | **None** | No relevant IP identified. | |
| | *Cry1C* | Gene of Interest | **None** | **None** | **None** | Use of *Cry1B* and *Cry1C* gene sequences fall under the patent family "Novel Genes Encoding Insecticidal Proteins" assigned to BASF Agricultural Solutions Seed (formerly Bayer Bioscience). These include chimeric gene constructs with *Cry1C* and dual Bt constructs with *Cry1B* . The scope of the US and Australian claims was narrowed during prosecution and are now limited to sequences that contain at least 98% homology to the claimed *Cry* DNA sequences. As the *Cry1B*$^{M}$ and *Cry1C*$^{M}$ DNA sequences have less than 98% DNA sequence identity to the *Cry1C* and *Cry1B* coding sequence covered by the BASF patent family, it is likely that use of these *Cry1*$^{M}$ sequences fall outside the scope of these patents. AU2007228981, EP1999141 and WO2007107302 applications were filed March 16th, 2007 and expire March 16th 2027.<br><br>No IP was found in the Indian online patent databases.<br><br>Note: We highly recommend that before using the *Cry1*$^{M}$ sequences, legal counsel be consulted to ensure that the sequences fall outside the scope of the BASF patent within the juridiction of interest. | AR059995; AT511515; AU2007228981; CA2646471; CN101405296; DK1999141; EA200802018; EP1999141; NZ571952; US20100235951; WO2007107302 |
| | *ME* 3'UTR | Transcription Regulation, 3'UTR | **Expired** | **Expired** | **None** | Use of the *ME* 3' UTR falls under the patent family "Plant transcription regulators from circovirus" assigned to CSIRO. An alignment using GenomeQuest shows there is 97% coverage and 99% identity between the *ME* 3'UTR used in constructs pJG1024/1027 and the CSIRO US 6211431 Seq ID 8. The claims also specify driving heterologous genes for insect resistance, in particular Bt toxins. These claims cover the proposed use SCSV promoters and *ME* 3' UTR in constructs pJG1024/1027. The Australian patent AU689311 was filed on August 30th 1995 and expired August 30th 2015. Likewise, the US patent US6211431 expired August 30th, 2015. No IP was found in the Indian online patent databases. | US6211431, AT266734T, AU689311b2, CA2198723A1, CA2198723C, CN1164869A, DE69533037D1, DE69533037T2, EP785999B1, ES2220935T3, JP10505233A, MX199701601A, Z291734A, WO1996006932A1 |
| **BT *Cry1B* DNA Cassette** | SCSV *S4* or *S4S4* | Promoter | **Expired** | **Expired** | **None** | As above | |
| | Tapetum *E1* Leader | Transcription Regulation, 5'UTR leader | **None** | **None** | **None** | As above | |
| | *Cry1B* | Gene of Interest | **None** | **None** | **None** | As above | |
| | Pea Rubisco *E9* | Promoter | **None** | **None** | **None** | No relevant IP was identified. | |

Note: Patents are territorial in nature. IP searhces uses a wide range of available on-line patent data to search by jurisdiction. However, where patents exist that are not digitally available, this research will not detect them. More in depth investigation requires hiring an agent in-country to search the local patent office. This may be particularly the case for India.
